# CurveCurator: A recalibrated F-statistic to assess, classify, and explore significance of dose-response curves

**DOI:** 10.1101/2023.08.07.552260

**Authors:** Florian P. Bayer, Manuel Gander, Bernhard Kuster, Matthew The

## Abstract

Dose-response curves are key metrics in pharmacology and biology to assess phenotypic or molecular actions of bioactive compounds in a quantitative fashion. Yet, it is often unclear whether or not a response significantly differs from a curve without regulation, particularly in high-throughput applications or unstable assays. Treating potency and effect size estimates from random and true curves with the same level of confidence can lead to incorrect hypotheses and issues in training machine learning models. Here, we present CurveCurator, an open-source software and interactive dashboard that provides reliable dose-response characteristics by computing p-values and false discovery rates based on a recalibrated F-statistic and a novel thresholding procedure. Application of CurveCurator to large-scale data sets demonstrates its scalable utility across several application areas.

## Introduction

Dose-response analyses are broadly applied in research from drug discovery and pharmacology to toxicology, environmental science, and epidemiology, to name a few. Prominent recent large-scale examples include phenotypic cell viability screens ^1–3^, activity-, affinity- or thermal stability-based drug-target binding assays ^4–6^, and proteome-wide drug-response profiling of post-translational modifications ^7^. Any dose-response experiment quantifies the response variable as a function of the applied dose and yields two orthogonal parameters: effect potency – the concentration producing the half-maximal response –, and effect size – the magnitude and direction of the response (Fig. 1a). Determining these parameters from dose-response curves is of immense practical relevance as this can e.g. guide drug discovery and repurposing, find the right dose for patients, define safety thresholds, and be used to train machine learning models for e.g. drug response prediction ^8^.

**Figure 1.**
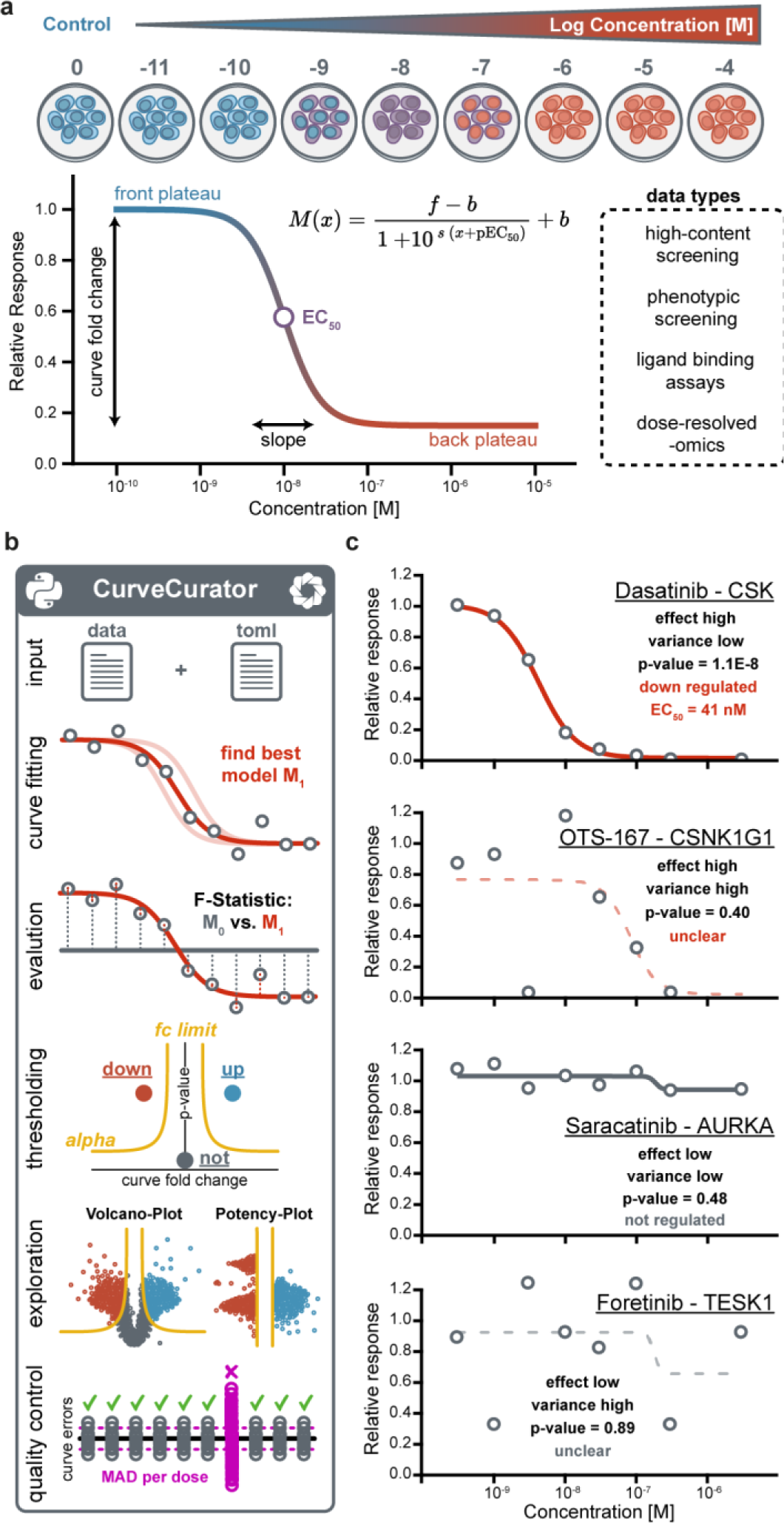
The CurveCurator pipeline. **a)** Dose-response curves can be described by characteristic curve parameters: front plateau, back plateau, slope, EC_50_, and fold change. These values are obtained by fitting a log-logistic model to data that can be of different types. **b)** The CurveCurator pipeline consists of python modules that are executed consecutively: The user provides a data and parameter file. CurveCurator will find the log-logistic model (M_1_, red line) that best describes the measured data points. Next, CurveCurator estimates a p-value for each curve by applying a recalibrated F-statistic, which compares the null curve M_0_ (no response, gray line) against the best curve M_1_ (red line). Curves are then classified into regulated, not-regulated, or unclear based on the significance asymptote alpha and the fold change asymptote. Results can be explored in an interactive dashboard that provides different views of the data. Optionally, a quality control analysis can be performed to identify experiments with high error across all curves. **c)** Curves from the Klaeger et al. data set exemplifying cases that need to be differentiated by CurveCurator. Only the EC_50_ values of significant curves reflect real potencies.

Several software tools exist that fit dose-response models and estimate potency and effect size ^9–14^. Surprisingly, none of them addresses the obvious question of whether or not the result constitutes a significant regulation. This is especially relevant for applications in which i) only a small proportion of cases produces significant regulations, ii) the effect size is close to the measurement variance of the assay, or iii) the assay is generally not very stable. So far, this has required (semi-)manual data evaluation by experts ^4–7, 15^, which suffers from inconsistent assessments among individuals and does not scale to the thousands to millions of curves generated by the aforementioned projects. Fitting a dose-response curve fit is, in essence, a regression problem. Hence, we propose that the statistical significance of dose-response curves can be assessed using F-statistics. Specifically, we evaluate the significance with respect to the null hypothesis that the response remains constant across doses and any observed variations in the response are due to random error. Related approaches have been used for assessing significance in the more general setting of time-series data, where the response is not expected to follow a sigmoidal shape ^16^ or the more specific setting of thermal stability as a function of ligand concentration, where temperature acts as the primary dimension ^17^. However, application of F-statistics to dose-response curves has not been extensively explored in prior publications. Here, we introduce CurveCurator, which estimates p-values from a recalibrated F-statistic and uses these to classify significant and non-significant curve regulation with low error rates.

## Main

CurveCurator is a statistics software pipeline for high-throughput dose-response data analysis (Fig. 1b). It is implemented as a Python package that is simple to install and use for people with little programming experience. Moreover, the package architecture enables integration of CurveCurator into existing pipelines. Multiple steps are parallelized, allowing large-scale datasets to be processed in a matter of minutes. To execute the pipeline, users need to provide response data (multiple input formats supported) and a simple parameter file in TOML format. CurveCurator fits a log-logistic model with up to four parameters (pEC_50_, slope, front, and back plateau; Fig. 1a) to the dose-response values. However, direct application of the F-statistic to dose-response curves yields poorly calibrated p-values (Fig. S1). The first issue is that the optimization surface for sigmoidal curves is non-convex (Fig. S2) so that curve fitting algorithms often get stuck in local minima leading to suboptimal F-statistic values compared to the fit at the global minimum. CurveCurator uses a novel heuristic that reaches the global minimum in almost all cases (Fig. S3).

The second issue is that the F-distribution with the corresponding degrees of freedom is not exact for non-linear models ^18^. Using randomly simulated null-hypothesis curves, a simple relationship was derived to calculate “effective” degrees of freedom that lead to properly calibrated p-values (Fig. S4ab). The validity of this approach was confirmed by producing uniformly distributed p-values in simulations and in real experiments, in which the vast majority of curves were expected to be unresponsive (Fig. S4cd). Next, CurveCurator classifies curves into three categories: significantly up-, significantly down-, and not-regulated. Classification is achieved by a method inspired by the fudge factor (s_0_) approach, also known as the Significance Analysis of Microarrays test (SAM) ^19, 20^. Based on user-specified asymptotes for p-value and effect size, a hyperbolic decision boundary is formed in a volcano plot, separating curves by statistical significance along the p-value axis and by presumed biological relevance along the effect size axis (Fig. S5). Rather than misusing the boundary to adjust the p-values ^21^, CurveCurator estimates the false discovery rate (FDR) of the user’s decision boundary using a target-decoy approach ^22, 23^. This can be done by simulating null hypothesis curves using the variance distribution of the input data and processing them with CurveCurator identically. The number of simulated curves above the decision boundary provides a false positive estimate. Using this approach, the FDR for curve classification can be controlled at the user’s preferred thresholds (Fig. S6). Comparing this approach to the Benjamini-Hochberg multiple testing procedure ^24^, which does not take effect size into account, showed that CurveCurator retains more regulated curves at low FDRs (Fig. S7). Finally, CurveCurator returns the fits and classifications in an output file with a similar format as the input file. Additionally, an interactive HTML dashboard enables data exploration locally in the browser of any desktop computer. We consciously chose <u>not</u> to classify curves that are unclear or otherwise noisy in order to obtain highly confident positive and negative groups of curves, which may be used as input for machine learning models in the future.

To demonstrate its utility, CurveCurator was applied to three dose-response data sets of different size and proportions of regulated curves. They illustrate how drug-phenotype, drug-target binding, and drug-PTM response data can be linked to elucidate the mode of action(s) (MoAs) of drugs, exemplified by the EGFR inhibitor Afatinib. First, the “target landscape of clinical kinase inhibitors” ^4^ was reprocessed (54,223 dose-response curves; 9% down-regulated), and drug-target classifications obtained by CurveCurator showed 97% consistency with the original manual analysis. Out of 247 assayed kinases, Afatinib was found to have nine significant interactions. As expected, the designated target, EGFR, was the most potent, followed by MAPK14 with a ∼100x lower potency (Fig. 2a). Second, the phenotypic CTRP cell viability screen ^3^ (379,324 dose-response curves: 63% down-regulated, 25% not regulated) was reprocessed (Fig. 2b). For Afatinib, 755 out of 760 cell lines were classified as significantly down-regulated, though the majority showed very weak potency. Due to the consistent potency dimension provided by CurveCurator, the above drug-target binding data and the viability data were put into context. This allowed a rough classification of cell lines into drug sensitivity groups driven by different targets of Afatinib (e.g., HER-, p38 MAPK-, and others). Of note, the plasma concentration of Afatinib in approved cancer therapies ^25^ precisely matches the border between HER-sensitive cells and p38 MAPK-induced cell toxicity, highlighting the importance of understanding the molecular MoAs behind dose-response relationships. Third, to understand the impact of Afatinib-target engagement on PTM-mediated signaling pathways, the decryptM profiles ^7^ of the EGFR-driven carcinoma cell line A431 were reprocessed by CurveCurator (19,596 dose-response curves: 5% down-regulated, 1% up- regulated, 46% not regulated; Fig. 2c). It is apparent that EGFR inhibition downregulates the direct EGFR substrate GAB1 pY627 at the expected potency. Downstream of GAB1, the MAPK- and AKT signaling axis were also inhibited with similar potencies (indicated by MAPK pT185/pY187 and AKT1S1 pT246) as were multiple transcription factors influencing cell growth such as ETV3, ELK1, and FOS (curves not shown). In contrast, inhibition of MAPK14 activity (monitored by MAPK14 pT180/pY182) only occurred at much higher drug doses as expected from Kinobeads. Excitingly, when combining the three orthogonal sources of information, the full cellular MoAs of Afatinib is revealed: Afatinib binds to EGFR with a K_D_ of ∼10 nM in cells (50% target engagement), inhibiting the main driver of this carcinoma and leading to the shut-down of key downstream survival signals. Substantial reduction in cell growth requires the inhibition of the bulk of signaling, which only occurs at ∼100 nM (90% target engagement). The observed inhibition of MAPK14 signaling at high drug doses is not relevant to the observed phenotypic effect on cell viability.

**Figure 2.**
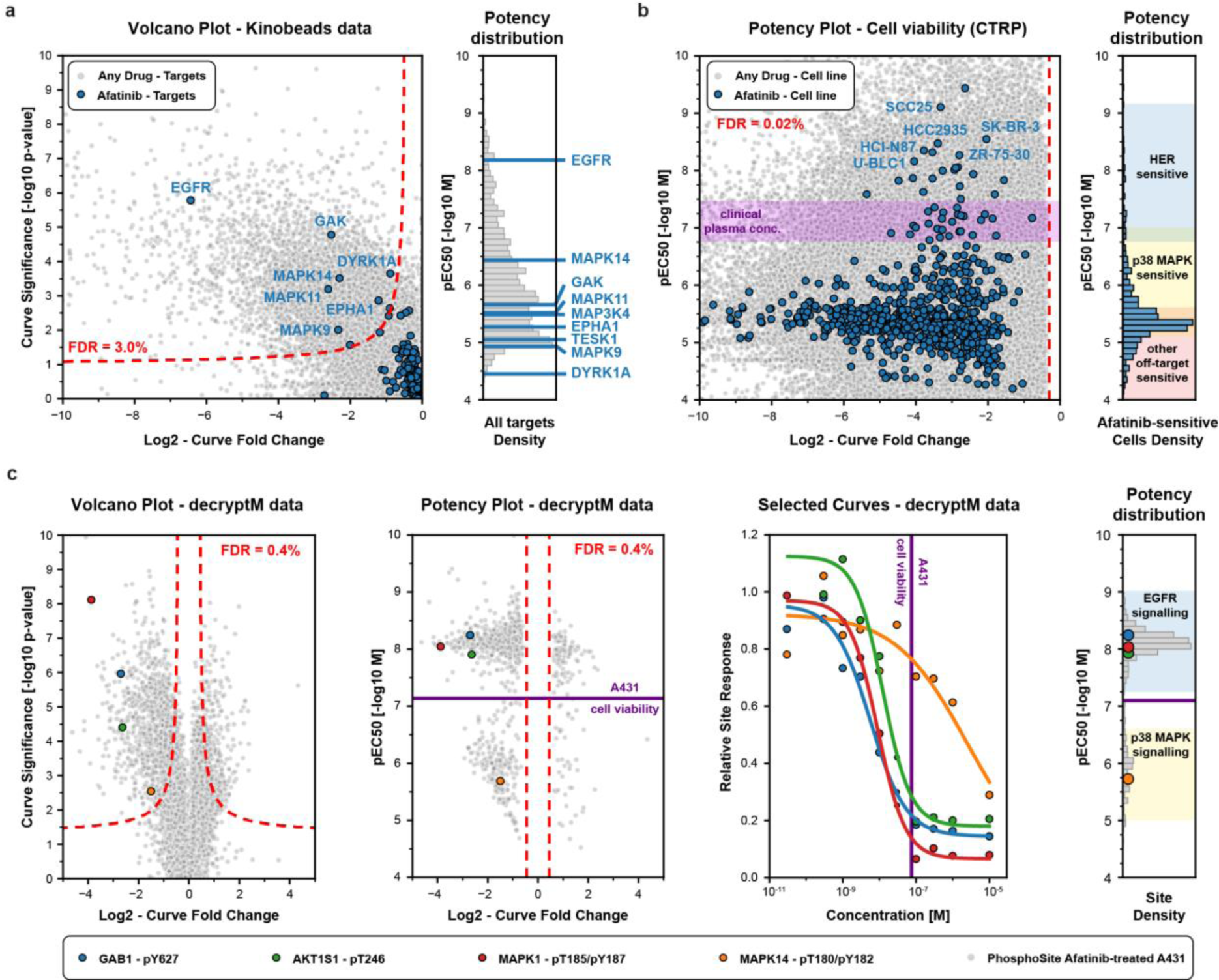
Application of CurveCurator to three data types relevant for drug MoA studies. **a)** Kinobead drug-target binding data. Significant drug-target interactions of the entire data set were identified using the CurveCurator pipeline and visualized in a volcano plot (left panel; significance vs. effect size; gray dots). The chosen decision boundary (red dashed line) obtained a FDR of 3.0%. Afatinib-target interactions are highlighted in blue, and significant Afatinib targets are labeled with their gene names. The pEC_50_ distribution in the right panel indicates the potency of Afatinib-target binding, highlighting EGFR as the most potent target. **b)** CTRP drug-cell viability data. Significant phenotypic responses of cell lines in the entire data set are marked in gray and Afatinib responsive cell lines in blue (left panel). Based on the potency-dimension, cell lines can be grouped into HER-, p38 MAPK-, or other target-sensitive (right panel). The purple band indicates the plasma concentrations achievable in Afatinib-based cancer therapies at the maximum tolerated dose. **c)** Cellular decryptM data of Afatinib-treated and EGFR-driven A431 cells. Significant phosphorylation responses in the entire data set are marked in gray and four example phosphorylation sites are shown in color (two left panels). The dose-response curves for the same four examples are shown in the second from right panel. The pEC_50_ distribution of all significant phosphorylation sites are shown in the right panel and can be grouped into EGFR- and p38-MAPK dependent signaling. The purple line indicates the potency of Afatinib in the phenotypic drug response assay.

## Summary

CurveCurator is the first tool that provides reliable p-values for assessing the statistical likelihood of regulation in dose-response experiments. It provides a simple means to classify curves and enables their exploration even in large-scale datasets. The classification of significant regulation combined with the use of pEC_50_ values allows the direct integration of multiple, complementary datasets by harnessing the perhaps most important characteristic of a drug in pharmacology - its potency.

## Methods

### Installation, Requirements, and Code Maintenance

CurveCurator is an executable, open-source Python module that can be downloaded from GitHub and PyPI. It can run on any operating system with an installed Python interpreter and a few common open-source libraries (numpy, pandas, scipy, statsmodels, bokeh, toml, tqdm). The easiest way to set up the correct environment is to use the package management and environment management system conda. We provide a step-by-step installation guide on GitHub as well as a detailed description of how to execute the script with different options. Furthermore, CurveCurator can be used as a library and thus be integrated into various existing analysis pipelines. Most of the source code is unit- and integration-tested to ensure stability and increase robustness for future updates and community collaborations. While the code will be maintained by the authors continuously, we encourage other researchers to contribute to CurveCurator for further standardizing dose-response analyses.

### Input and Output

To execute CurveCurator, the user must only provide the data file with dose responses and a simple parameter file in TOML notation. In the most generic case, any raw data can be provided as a simple tab-separated file containing the response observations for all doses as well as a sample identifier column. Additionally, CurveCurator can directly parse proteomic data on protein- or peptide-level coming from various search engine outputs, including MaxQuant, ProteomeDiscoverer, and DIA-NN. The TOML parameter file contains all relevant information to interpret the data and run the pipeline. Additionally, the user can add optional parameters to adapt the pipeline to the requirements of a specific experiment and data type. We recommend storing this TOML parameter file next to the input and output files to increase reproducibility and transparency. All output files of CurveCurator will be stored relative to this TOML file by default. A log file reports each step, pipeline values, overall processing times, and errors if occurring. Detailed explanations, as well as example parameter files, are available on GitHub.

The CurveCurator pipeline returns multiple files, including the curves.txt file. It contains relevant columns from the input data together with curve fits, statistics, and classification columns. The dashboard.html file generated by bokeh enables interactive exploration of the curve data in the browser on any computer. Selected curves can be exported as figures or data tables. If the user has normalization activated, the applied log_2_-transformed normalization factors are reported. If systematic biases are recognized, we encourage the user to drop the affected dose for all curves in the data set and re-run the experiment.

### Data pre-processing

CurveCurator will first aggregate duplicates in the provided raw data. For proteomics data, this will result in unique proteins or modified peptides depending on the data type. Missing values can be included, imputed, or filtered as specified by the user in the TOML file. We do not recommend imputing missing values in viability data sets as these values are typically missing at random and are difficult to estimate. For proteomics data, missing values are typically missing not at random but are the consequence of low-intensity sampling bias and thus can often be imputed with a low-intensity number. The imputation value is drawn from the intensity distribution of the data. The user can specify the desired quantile. Next, intensity values can be globally normalized. This is again an option mostly relevant for proteomics data, where typically the protein distributions stay constant over the course of the treatment and up- and down-regulation balance or do not make a significant proportion (>50%) in the data set. If the user wants to apply more advanced normalization procedures, it is always possible to do this prior to the CurveCurator pipeline and skip the aforementioned standard procedures. Finally, the processed data is then transformed into ratios relative to the control sample.

### Finding the best model

CurveCurator uses two competing models, which are evaluated based on the observed responses (normalized to the control sample, see above). To obtain model parameters, CurveCurator currently supports ordinary least squares (OLS) regression as well as maximum likelihood estimation (MLE). While the mean model has an analytical solution, it does not exist for the log-logistic model and thus requires minimization procedures of the OLS cost or MLE negative log-likelihood objective functions, respectively.

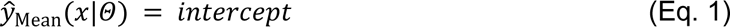

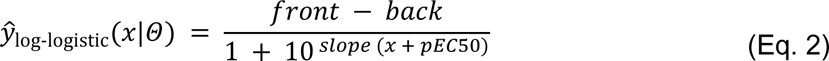

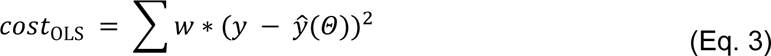

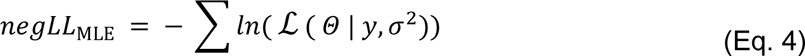

Because the minimization surface of the sigmoidal curve is non-convex (a surface with many local minima) for the slope and pEC_50_ parameter, there is no guarantee to find the global minimum from all start points (Fig. S2). CurveCurator provides different optimization strategies to overcome local minima, including different start points and multiple minimization rounds. This is, however, always a compromise between the global success rate and the overall run time (Fig. S3). The user can choose from the following four minimization strategies through the speed parameter in the TOML file: in “fast” mode, a sequence of pre-defined guesses (calculating pEC_50_, front, back based on the data) is evaluated, and only the best initial guess is then minimized; in the “standard” mode, the same best initial guess is evaluated, and then multiple minimizations are performed using different slope parameters together with the best initial parameters; in the “extensive” mode, the entire series of alternative start guesses is directly minimized; and in the basin-hopping-mode, the global minimization algorithm “Basin-Hopping” is applied to overcome local minima. If multiple optimizations are performed per curve, the best solution is taken and assumed to be the global minimum for statistical analysis. All minimizations are bounded (pEC_50_: experimental drug range +-4 orders of magnitude; slope: 0.01 - 20, front & back: 1e-3 - 1e6). MLE uses the Nelder-Mead minimization algorithm, and OLS uses the L-BFGS-B algorithm supplemented with the Jacobian matrix to speed up the minimization. CurveCurator supports weights in the OLS approach that can be specified by the user for each data point, which can make curve estimation in some scenarios more robust. A subset of the model parameters can also be fixed to a pre-defined value if the user’s experimental setup allows for this assumption. The optional interpolation method can create more robust fits by linearly interpolating helper points between data points during the fitting procedure.

### F-statistics and p-values

The best log-logistic model from the fitting procedure is then statistically evaluated for significance, and only significant log-logistic models should be interpreted with regard to the fitted values, e.g. pEC_50_ value.

For linear models, the p-value procedure is well-known and widely applied. The basic idea behind the F-value in regression problems is to quantify how much better a more complex model (M_1_ with *k* parameters) fits the data compared to a simpler model (M_0_ with *j* parameters and *j*<*k*) given the *n* observed data points and the corresponding sum-squared errors (SSE).

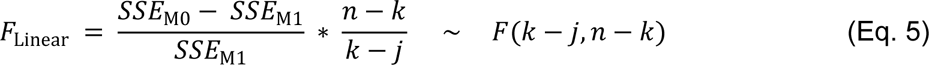

Although not a linear model by nature, the log-logistic function still meets the required assumptions of random sampling, independence of observations, residual normality, and equal variance of the errors. The basic rationality behind CurveCurator’s recalibrated F-statistic is similar to the linear F-statistic. It also quantifies how much better the fitted log-logistic model (M_1_) is compared to the mean model (M_0_), which describes that there is no relationship between the applied dose and the observed response. We found, however, that n/k was a more appropriate scaling factor for the 4-parameter log-logistic function.

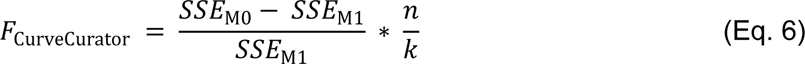

The obtained F-value can then be used to calculate a p-value that quantifies how often a curve with a similar or bigger F-value can be found by random chance. We observed that these F-values follow a parameterized F-distribution with degrees of freedom that diverged from the case of linear models. Using extensive simulations under the null hypothesis, we obtained a simple function to calculate the “effective“ degrees of freedom as a function of *n*. A small correction was additionally necessary for curves with low number of *n* data points to follow the approximated F-distributions, potentially indicating over-fitting of the log-logistic model when too few observations are present for 4 parameters (Fig. S4). The location parameter of the F-distribution was found to be 0.12 for all *n*, indicating that dose-response curves with two flexible plateaus can always remove more error than the intercept model M_0_.

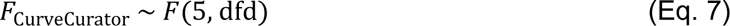

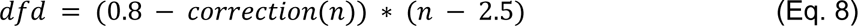

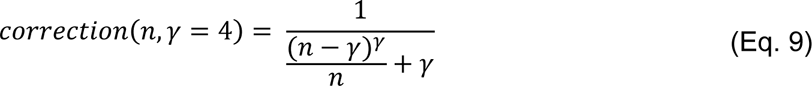

### Thresholding and false positives

Each curve has a p-value that can be directly interpreted. If only the p-value is considered to identify significant curves, multiple testing corrections, e.g. Benjamini-Hochberg procedure, can directly be applied to control for false positives in large datasets (Fig. S5). This procedure can be activated via the TOML file, and CurveCurator supports different multiple testing procedures. However, the (default) recommended thresholding procedure uses both the p-value and curve fold change to identify significant curves. The curve fold change is defined as the log_2_-ratio between the lowest and highest concentration using the regressed model.

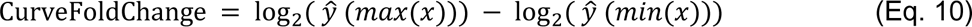

First introduced as the SAM-test for differential analyses ^19^, the combination of these two dimensions leads to a hyperbolic decision boundary in the volcano plot that separates hits by statistical significance and biological relevance simultaneously. We transferred this SAM principle of differential T-statistics to the curve F-statistic to obtain equivalent decision boundaries for the dose-response curve analyses. This was possible by recognizing that a dose-response curve converges in the limit to two groups (front plateau group = not affected data points, and back plateau group = affected data points), where the curve fold change is equivalent to a conventional SAM fold change between the two plateau groups. In this case, *F* = *T*^2^ allowing for the conversion and application of the s_0_ SAM principle. CurveCurator simplified this process by calculating the tuning parameter s_0_, however, directly from the user-specified significance and fold change asymptotes. The user only needs to specify an alpha asymptote, which is the p-value threshold that a curve needs to pass no matter how big the effect is, and a fold change asymptote, which specifies the minimal required effect no matter how significant a curve is.

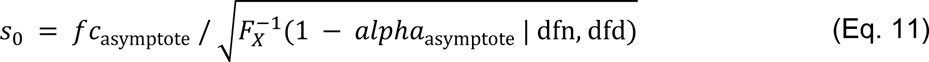

Where 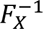 (*x* | dfn, dfd) is the inverse cumulative density function of an F-distribution with degrees of freedom *dfn* and *dfd* as determined in the section above (Fig. S5).

Applying a hyperbolic decision boundary is also an efficient way to reduce the FPR (false positive rate) because highly significant yet random curves tend to have small effect sizes. Consequently, the same FPR can be achieved by various combinations of the two asymptotes (Fig. S5). This phenomenon must be considered and has substantial implications for multiple testing corrections because often, the fold change asymptote reduces the FPR to such a small number that the total number of expected false positives in real-world applications is negligible. This can also be seen when the same tuning parameter s_0_ that defines the hyperbolic decision boundaries is used instead to transform the curve’s F-value into the s_0_-adjusted F-value (*F_adj,_*_i_) based on s_0_ and the curve’s measured fold change (*fc_i_*).

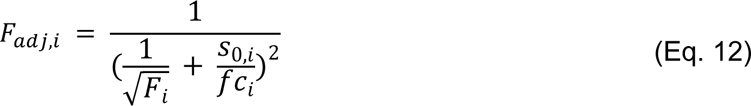

We then transform this s_0_-adjusted F-value into a *”relevance score”* using the cumulative density function *F_X_ ( x | dfn, dfd)*. For s_0_=0.0, this simply corresponds to the p-value of the curve. For s_0_>0.0, this can no longer be interpreted as a p-value ^21^, but it still provides an appropriate ordering of curves by both statistical and biological relevance. Additionally, a - log_10_ transformation is applied for visualization purposes.

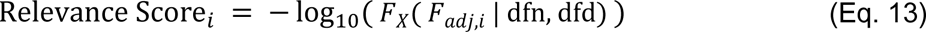

After these two transformation steps, the previous hyperbolic decision boundary becomes a horizontal decision boundary in the volcano plot of fold change vs. relevance score (Fig. S6).

Next, the FDR (false discovery rate) can be estimated using a target-decoy approach. Decoy curves are generated based on the sample variance distribution (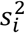) of the input data directly. From this variance distribution, a variance value is randomly drawn for each decoy curve and the basis for a random curve generator assuming the null-hypothesis is true.

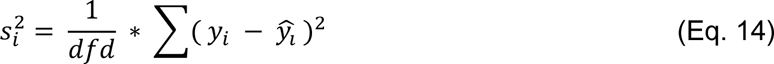

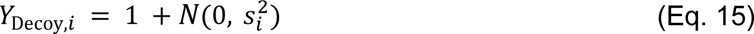

The decoy curves are then subject to the identical analysis pipeline. Finally, the target-decoy relevance score distributions are compared allowing the calculation of a q-value for each curve and the overall FDR of the user’s pre-defined thresholds (Fig. S6).

Note that, one could technically adjust the FDR to a pre-defined threshold by moving the decision boundary up or down in the relevance score volcano plot. However, as the relevance volcano plot only remains unchanged for a fixed value of s_0_, this will automatically adjust both the alpha and fold change asymptotes simultaneously. Therefore, we explicitly recommend against finding a threshold based on a pre-given FDR-value, but instead encourage users to define the significance and fold change asymptotes first and accept the resulting FDR. Especially in datasets with a high proportion of regulated curves, e.g. viability data, increasing the FDR to 0.1% - 10% will lead to the inclusion of many undesirable curves. Users can activate the FDR analysis via the --fdr option.

### Not-Regulated Curves

For many applications, it is also useful to know which dose-responses show clear independence of the doses. It is fundamentally impossible to prove absence of an effect statistically. Thus, we developed a heuristic approach using the null model to classify only a clear non-responsive line to be not regulated. A clear non-responder has a mean-model intercept close to 1.0 and is allowed to maximally diverge +-fc_lim/2. Additionally, the variance around the null model should be low and is quantified by the root-mean-squared error (RMSE). By default the maximal tolerated variance is a RMSE of 0.1, but can be adjusted by the user. Optionally the user can add additional criteria such as a p-value threshold to be even more restrictive to the non-classification.

### MAD-Analysis

CurveCurator is able to detect systematic problems in data sets. This is particularly relevant for proteomics data, where one raw file or TMT channel can exhibit disproportionately high variance and interfere with the statistical evaluation. The median absolute deviation (MAD) analysis is a simple way of detecting aberrant variance by looking at residual distributions over all curves in the dataset. A MAD-value of, e.g., 0.1 means that 50% of all curves are further away from the model estimate than 0.1 ratio units. For proteomics data, we consider high-quality data to be <0.1 and exclude data points if 0.15 is exceeded. As most measurements are close to 1, these values can be interpreted as 10% and 15% variance thresholds. Users can activate the MAD analysis via the --mad option.

### Dashboard

CurveCurator outputs an interactive dashboard that allows for fast and interactive data exploration. All dashboards have a global overview plot on the left side, which can switch between volcano and potency representation. Each dot is one dose-response curve, and the color indicates its potency. The Volcano plot relates the curve fold change to the curve significance. The Potency plot relates the curve fold change to the curve potency measured by pEC_50_. The global plot is interactive, and the dose-response area plots the selected curves and yields a quick overview of the raw data and the fitted curve. More information about a selected curve can be found at the bottom table. Adjustable filters can help to focus on specific subparts, and search boxes can help to select and search for specific curves. Additional functionalities are available depending on the specific dataset and columns present in the input.

### External Datasets

*<u>Kinobeads</u>* ^4^ search results and LFQ intensities were downloaded from PRIDE (PXD005336) and filtered for direct binders (255 proteins that can bind to the Kinobeads via an ATP pocket including 216 kinases). The direct binder list, experimental design table, and the manual target annotations were obtained from the original publication. Binders with less than two data points per curve or missing in the control experiment were excluded. The remaining missing values were imputed per experiment using the 0.5% intensity quantile. The resulting Kinobeads matrix consists of 278 unique drugs with 9 data points each and was subjected to CurveCurator. For competition pulldown data, the alpha value was set to 10% and the fold change asymptote was set to 0.5.

*<u>CTRP Viability data</u>* ^3^ was downloaded using the *downloadPSet* function of the R package PharmacoGx. This consisted of 373,324 drug-cell line combinations from 545 drugs and 887 cell lines, each with n=16 doses plus one control. Because the CTRP screen featured many different dose ranges, the data was split into separate CurveCurator input files for each dose range. Note that most drugs were profiled at a single dose range. Missing values were very sparse and therefore there was no need for imputation. For CurveCurator analysis, the alpha value was set to 5% and the fold change asymptote was set to 0.3. Fold change was computed relative to the control instead of the lowest dose as several drugs showed significant regulation at the lowest dose already. The different dose-range output files were re-combined and the FDR was estimated once for all curves.

*<u>decryptM</u>* ^7^ search results and TMT intensities were downloaded from PRIDE (PXD037285) and the datasets ”Dasatinib Triplicates Phosphoproteome MS3” and “3 EGFR Inhibitors Phosphoproteome” were re-analyzed in this paper. Experiments that were searched together were separated and subjected to CurveCurator individually. The experimental design table was obtained from the original publication. For TMT peptide data, the TMT channels were median-centered and missing values were imputed. Peptides with more than 4 missing values were excluded. The alpha value was set to 5% and the fold change asymptote was set to 0.45. This fold change asymptote is 46% less stringent than in the original publication, which was solely possible due to the gained specificity of the hyperbolic decision boundary. Cell viability data for A431 was used from the original publication.

### Limitations

CurveCurator is looking specifically for dose-response shapes that can be described by the 4-parameter log-logistic model reflecting a single drug-target effect. While this assumption is valid for most experimental curves, there are cases where the dose-response curve is shaped by multiple effects resulting in a non-sigmoidal curve shape ^11^. The consequence is that CurveCurator cannot explain the true complex relationship well, typically resulting in big effect sizes but poor p-value. In the future, we envision that CurveCurator supports a wider variety of models, given that the p-values can be well-calibrated for those more complex models. Another limitation is that the fold change calculation requires the lowest dose close to the front plateau to get a good estimate. However, if a drug is more potent than the applied experimental dose range, the fold change estimate is compressed leading to an increased rate of false negatives among the most potent effects. To overcome this we recommend to either choose the experimental drug range more carefully after some pilot, or use a modified fold change calculation relative to the control at the cost of increased false positives.

## Data availability

CurveCurator output files for all datasets, as well as interactive dashboards are available on Zenodo https://doi.org/10.5281/zenodo.8215092

## Code availability

Software and documentation is freely available on GitHub under open-source license Apache 2.0. : https://github.com/kusterlab/curve_curator (Additionally, the accepted version of the code used in this paper will be permanently available on Zenodo with a stable doi). Example data set are also available on GitHub.

## Acknowledgements

The authors want to thank all members of the Kusterlab for critical but productive discussions. Special gratitude goes to all the internal and external beta users for their helpful feedback, which improved the tool in many instances.

## Author contributions

F.P.B. conceived the CurveCurator approach to calculate p-values and implement the SAM thresholding for curves. F.P.B., M.G., and M.T. wrote code and performed data analysis. B.K. and M.T. directed and supervised code and data analysis. F.P.B., B.K., and M.T. wrote the manuscript.

Correspondence to M.T.

## Supplementary information

**Fig S1.**
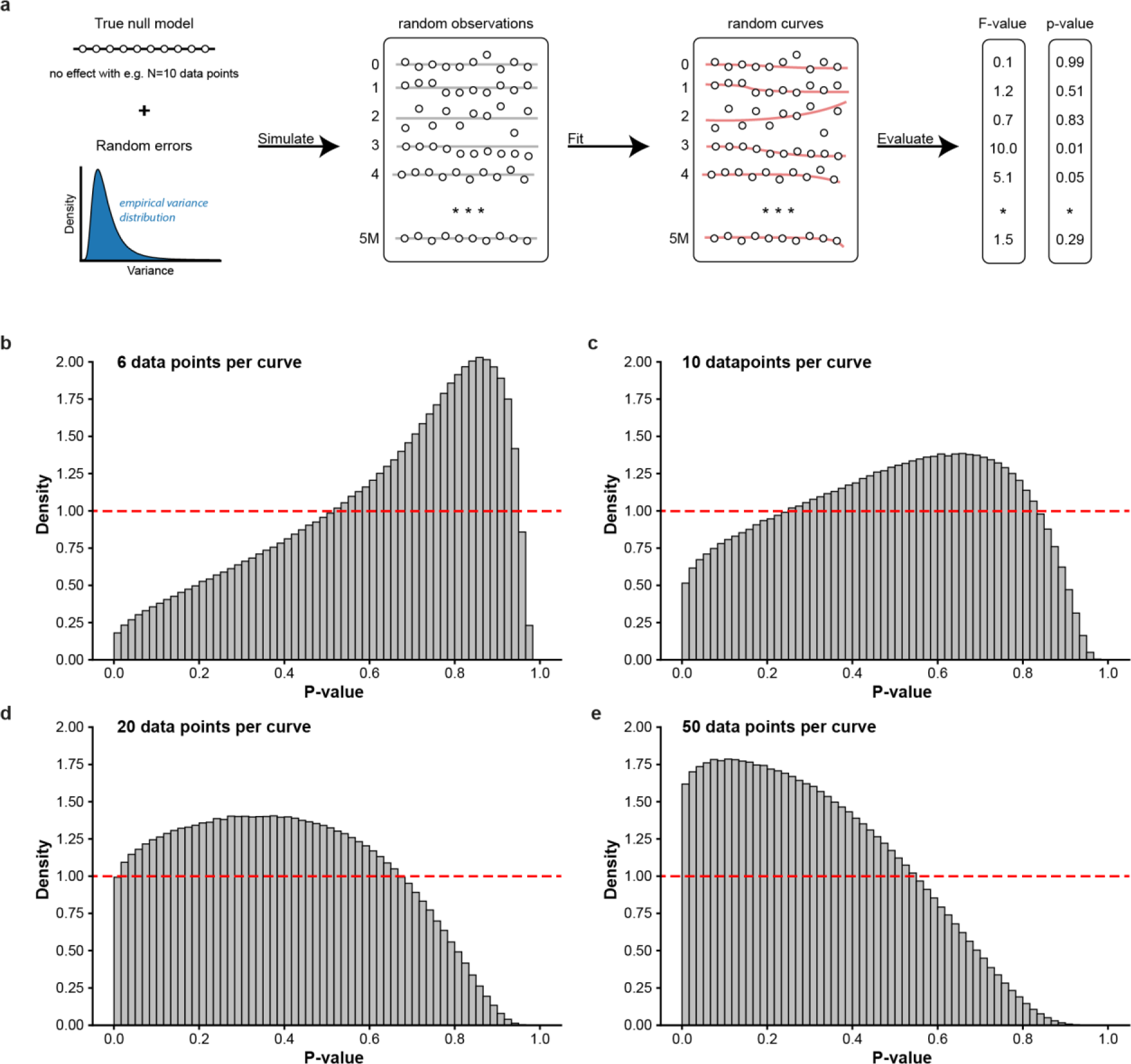
P-value distributions for curves under the null hypothesis are not well calibrated using the standard F-statistic. **a)** To analyze the statistical properties of dose-response curves, we simulated 5 million dose-response curves under the null hypothesis that the response is independent of the dose (no dose-dependent changes, intercept = 1.0). Random errors were introduced by first drawing a variance value from an empirical variance distribution for each null model and second adding a normally distributed random error (*N*(0, variance)) to each data point to obtain a simulated set of random observations (Equations 15). The 4-parameter log-logistic function (Equation 2) was then fitted to each simulated set of data points, and F- and p-values were obtained following the standard approach for linear models (Equation 5). We performed these simulations for **b)** n=6 doses, **c)** n=10 doses, **d)** n=20 doses, and **e)** n=50 doses. Well-calibrated procedures should produce a uniform p-value distribution with a density of 1.0 (red dashed lines in all panels). For simulated data with few doses (n=6 or n=10), p-values are generally too high (conservative). For simulated data with many doses (n=20 or n=50), p-values are generally too low (anti-conservative).

**Fig S2.**
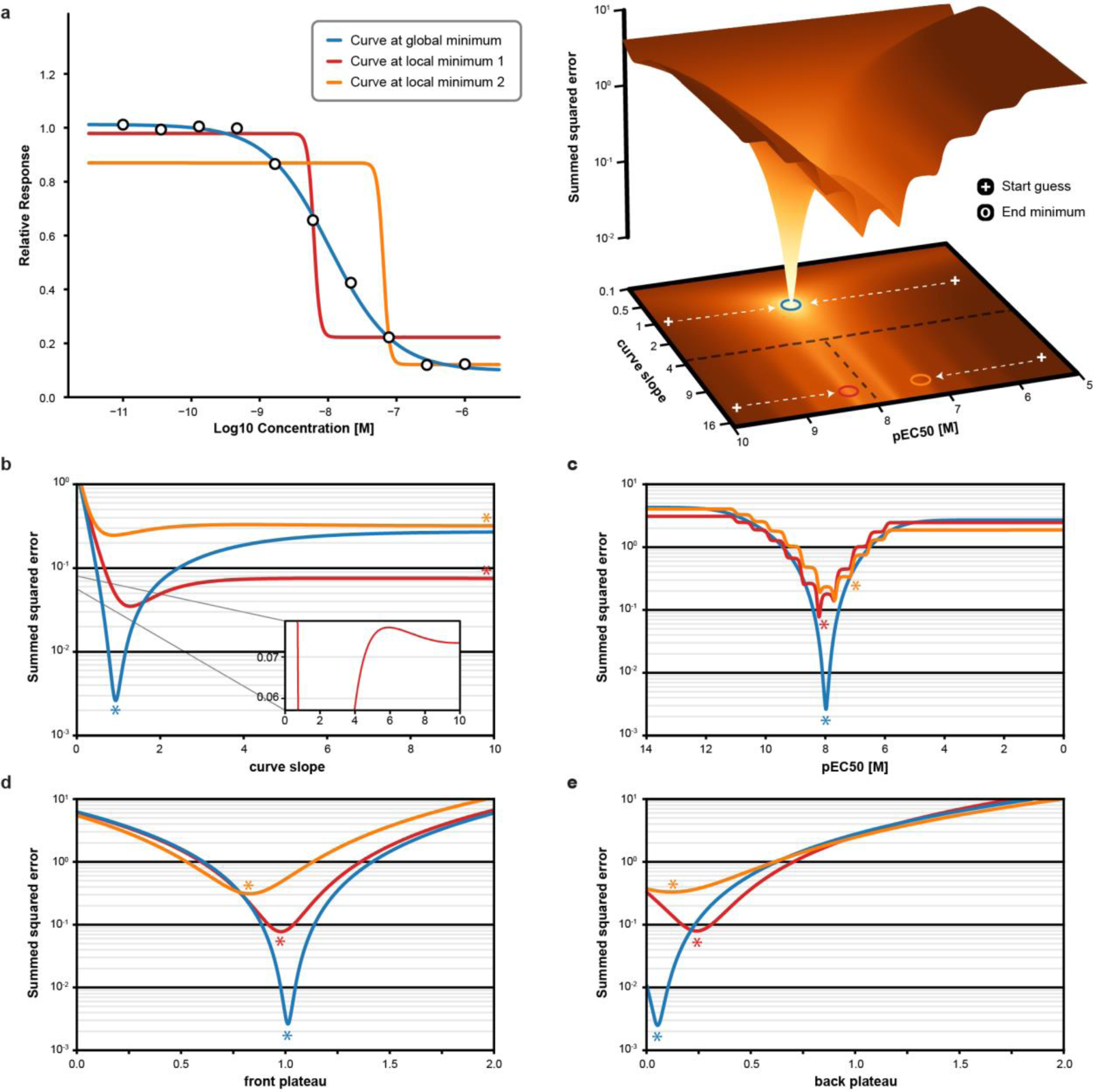
The minimization surface of the 4-parameter log-logistic function is not convex, leading to undesired convergence of curve fits to local minima. **a)** A dose-response curve was simulated (left panel; black data points, n=10) by adding random error to a down-going, ground-truth 4-parameter log-logistic model (pEC_50_=8.0, slope=1.0, front plateau=1.0, back plateau=0.1). Three different initial guesses for the four parameters were tested, leading to three different curve fits (red, orange, and blue). Each initial guess converged to a different minimum, of which only one converged to the global minimum (blue) to achieve the best possible fit. The cost surface of slope vs. pEC_50_ is obtained from ordinary least squares for the same simulated dose-response curve (right panel). The color from white to brown indicates the summed squared error at one particular slope-pEC_50_ pair (white: good fit; brown: poor fit). Initial guesses are represented as white pluses on the 2D projection, and the arrows indicate the simplified trajectories from those initial guesses to the next minima (circles) during the minimization process. Two initial guesses converge to local minima (red and orange circle), whereas the other initial guess converges to the global minimum (blue circle). **b-e)** One-dimensional cost plots based on the three minima (red, orange, and blue curves) from panel a when changing only one parameter at a time (slope, pEC_50_, front plateau, back plateau) while fixing the other curve parameters with the values of the respective minimum (indicated by the asterisks). It shows that for the slope and pEC_50_ parameters, the cost functions are non-convex (multiple minima), and thus, the curve fit algorithm can get trapped in these local minima. The front and back plateau parameters have convex cost functions instead and thus will always converge to the locally optimal parameter value for any combination of the fixed parameters.

**Fig S3.**
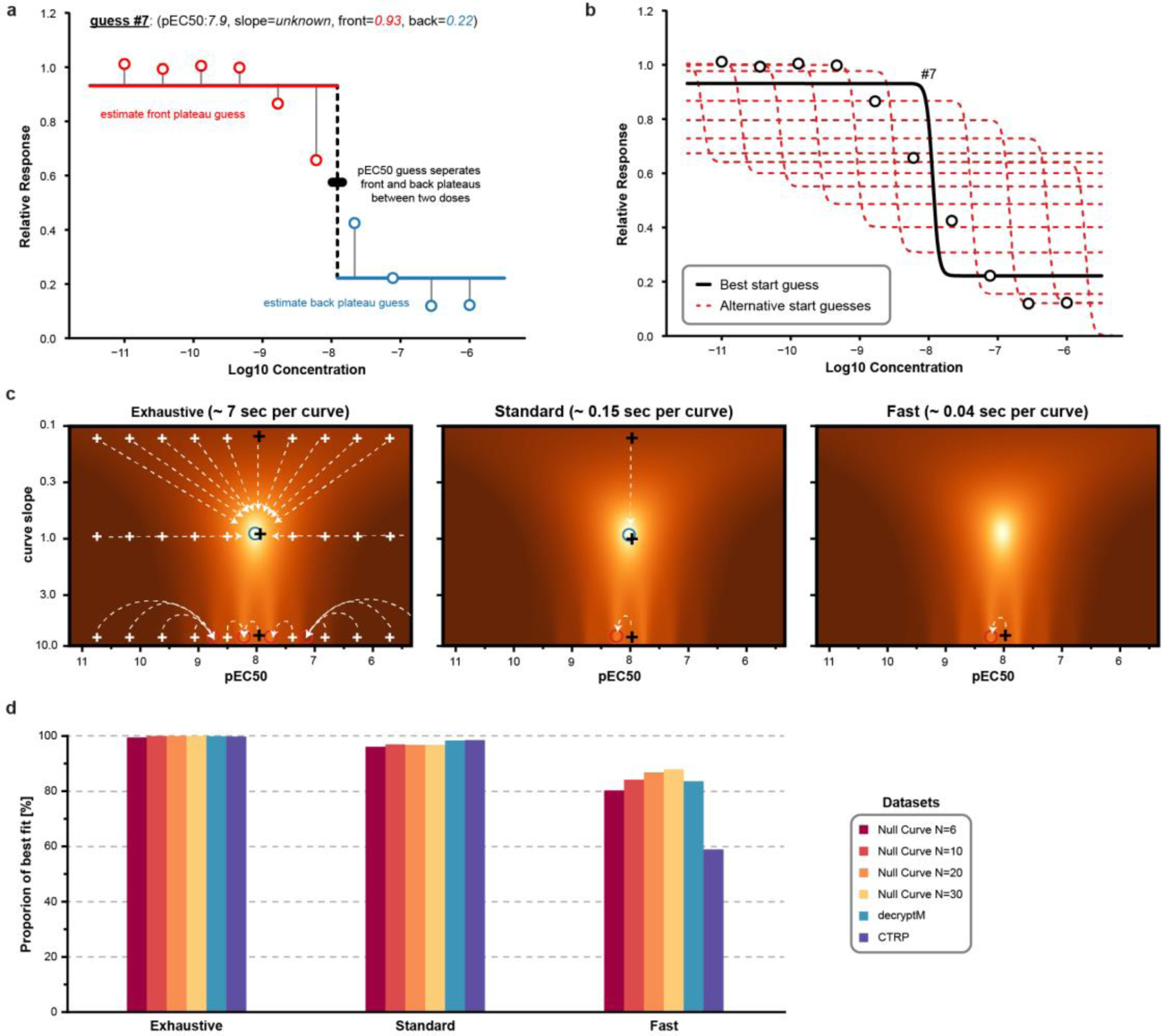
Three strategies for setting initial parameter guesses lead to different success rates in reaching the global minimum. **a)** CurveCurator generates multiple initial guesses calculated from the measured response values (circles). The guessing procedure is exemplified for the start guess #7. Here, the pEC_50_ value is between data points 6 and 7 at approximately 12.5 nM. The data points are then divided into a front (<pEC_50_, red) and back section (>pEC_50_, blue), and the mean is used to calculate the plateau guesses values, respectively. The slope is yet undetermined. **b)** The cost of each possible start guess (1 - 11, red dashed lines) is evaluated by ordinary least squares to determine the best guess (black solid line), which happens to be guess #7 from panel a. **c)** CurveCurator implemented three different minimization strategies (exhaustive, standard, fast) that use different numbers of start guesses to increase the odds of reaching the global minimum. The position of these start guesses on the cost surface is depicted with white crosses. The best start guesses are shown as black crosses. Note that the cost surface will look different for each dose-response curve. “Exhaustive” uses all alternative start guesses with three different slopes and is, consequently, the slowest method and scales with N doses. Many start guesses converge to the same few minima (dotted white lines). “Standard” is more efficient and uses only the best guess with three different slopes. “Fast” only uses the best initial guess with a single arbitrary high slope value. **d)** We evaluated how often the global minimum was reached by the three strategies (exhaustive, standard, fast) for different numbers of doses (n = 6, 10, 20, 30) in simulated null curves as well as in real data sets, e.g., decryptM (Dasatinib in K562) and CTRP cell viability data. The global minimum was defined as the best F-statistic with an error tolerance of 0.1%. The standard strategy turns out to be the best compromise between time and fitting success.

**Fig S4.**
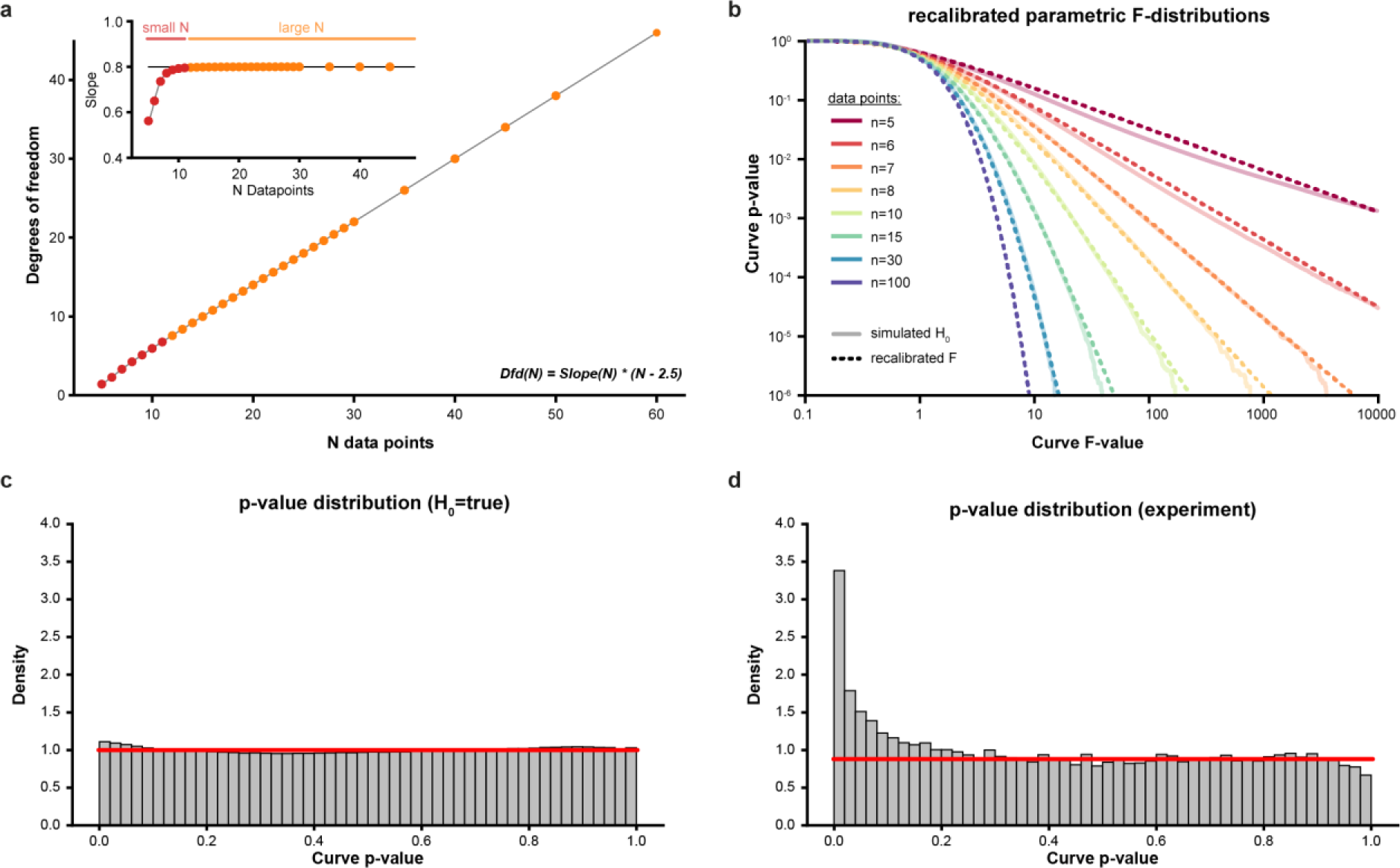
CurveCurator estimates effective degrees of freedom to obtain well-calibrated p-values. **a)** CurveCurator estimates the effective degrees of freedom (F-parameter dfd) based on the number of doses n in an experiment. This relationship is linear for large N (orange data points, black line, Equation 8). For small n (n<11, red data points), a penalty factor has to be introduced to account for overfitting (Equation 9). This relationship was determined from 5 million random simulations under the null hypothesis. **b)** log-log plots of F-values vs. p-values for different numbers of doses (different colors) based on 5 million random simulations. Using the relationship of panel a, the recalibrated parameterized F-distributions (dashed line) are able to approximate the real null distributions (solid line) (Equation 7). **c)** Uniform p-value distribution resulting from recalibrated F-distributions (5 million H_0_-simulations, n=10 doses). The red line indicates the expected density of 1.0. **d)** P-value distribution of experimental data (3 Dasatinib replicates in K562, n=10 data points, from Zecha et al.). The p-value distribution of the true negatives follows a uniform distribution. Curves above the red line are presumably truly regulated by Dasatinib.

**Fig S5.**
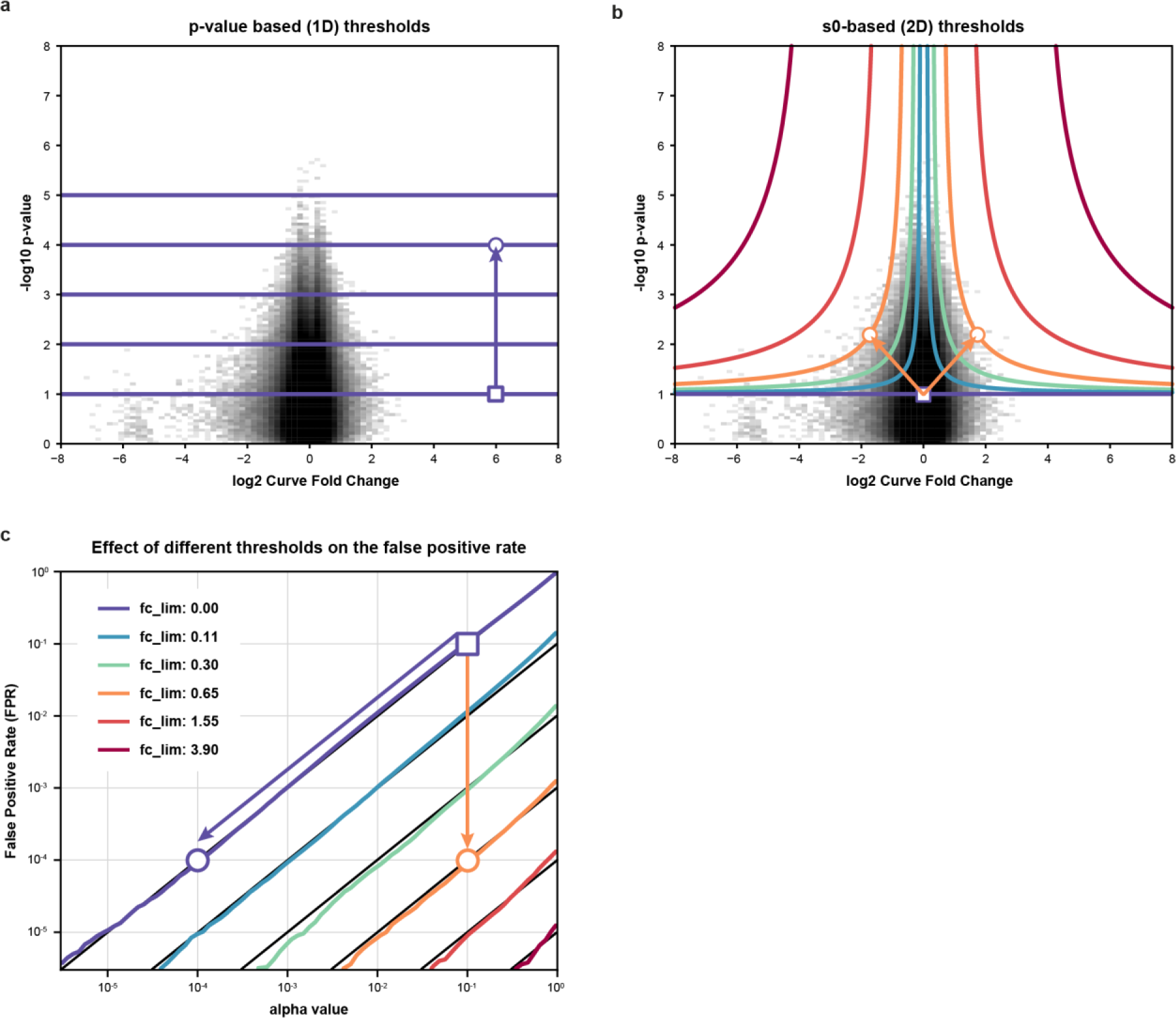
Determining the significance of dose-response curves using the s_0_ principle leads to acceptance of a greater number of biologically relevant curves at the same FPRs. **a)** Volcano plot (significance vs. effect size) of 5 million simulated null hypothesis dose-response curves (n=11 doses). Random curves with high significance tend to have small effect sizes. Moreover, real curves with small effect sizes are also typically not of (biological) interest because they are difficult to interpret. Traditional decision boundaries consider only the p-value (horizontal lines), where a user-defined alpha threshold (purple lines) establishes the false-positive rate (FPR). Lowering the FPR (box-to-circle arrow) can only be achieved by setting a more stringent alpha threshold, likely losing many true positives. **b)** Same volcano plot as in panel a, but showing several hyperbolic decision boundaries using the SAM s_0_ principle. These colored hyperbolic decision boundaries combine a particular fold change asymptote with a fixed significance asymptote of alpha=0.1 and reject curves with small effect sizes but high significance. For example, the orange decision boundary has the same FPR as a traditional alpha threshold of 1e-4 but would accept biologically relevant dose-response curves with e.g. a p-value of 1e-3 and a fold change of 2.0, which would have been rejected by the traditional alpha threshold. **c)** For a fixed value of the fold-change asymptote, there is a linear relationship between the value of the alpha asymptote and the observed FPR. Consequently, the same FPR can be achieved by multiple combinations of fold change and alpha asymptotes. The purple arrow indicates a reduction of the FPR to 1e-4 by reducing the alpha threshold, as depicted in panel a. The orange arrow indicates the same reduction of the FPR by introducing a fold-change asymptote but keeping the alpha asymptote at 0.1, as depicted in panel b.

**Fig S6.**
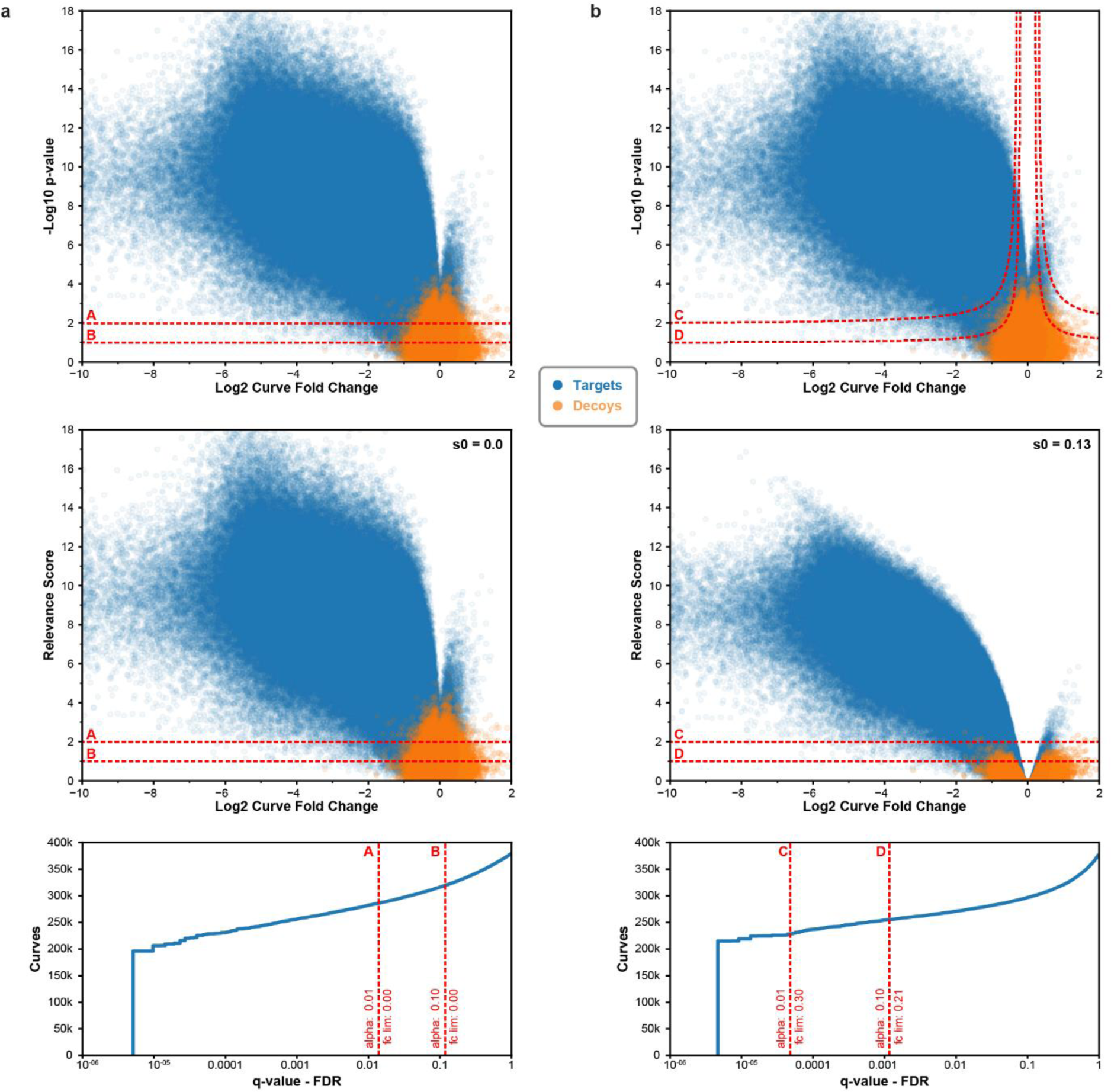
CurveCurator uses a target-decoy strategy to estimate the false discovery rate for the user-defined relevance asymptotes. **a)** The upper volcano plot shows the CTPR cell viability data set (blue) and its simulated decoys (orange) using the same measurement variance distribution as the experimental data. The two decision boundaries, A (alpha=1%) and B (alpha=10%) do not use a fold change asymptote (fc_lim = 0). Consequently, the fudge factor s_0_ is also 0.0, and the transformation of significance (upper panel y-axis: p-value) into the relevance score (middle panel y-axis) results in the same numerical values. Using the decoy distribution, the FDR of decision boundary A and B can be estimated (lower panel). Target and decoy curves with highly significant p-values but small fold changes pass the decision boundary A & B, resulting in FDRs of 1.4% and 11.8%, respectively. **b)** The upper volcano plot is identical to the one of panel a, but this time, two hyperbolic decision boundaries, C and D, are used to identify significant curves. The alpha and fold change limits of C and D were chosen such that they have the same transforming fudge factor (s_0_=0.13). Using s_0_, the statistical significance is transformed into the relevance score (Equations 12, 13), leading to a stronger reductive effect the smaller a curve fold change is. Similarly, this transformation bends the previous hyperbolic decision boundaries to horizontal decision boundaries. This results in a "relevance volcano plot" where the y-axis shows the curve’s relevance score instead of its p-value. Because many decoys have small fold changes, fewer decoys pass the relevance decision boundary C & D, resulting in much lower FDRs of 0.005% and 0.1%, respectively, although they have the same alpha value as A and B from panel a.

**Fig S7.**
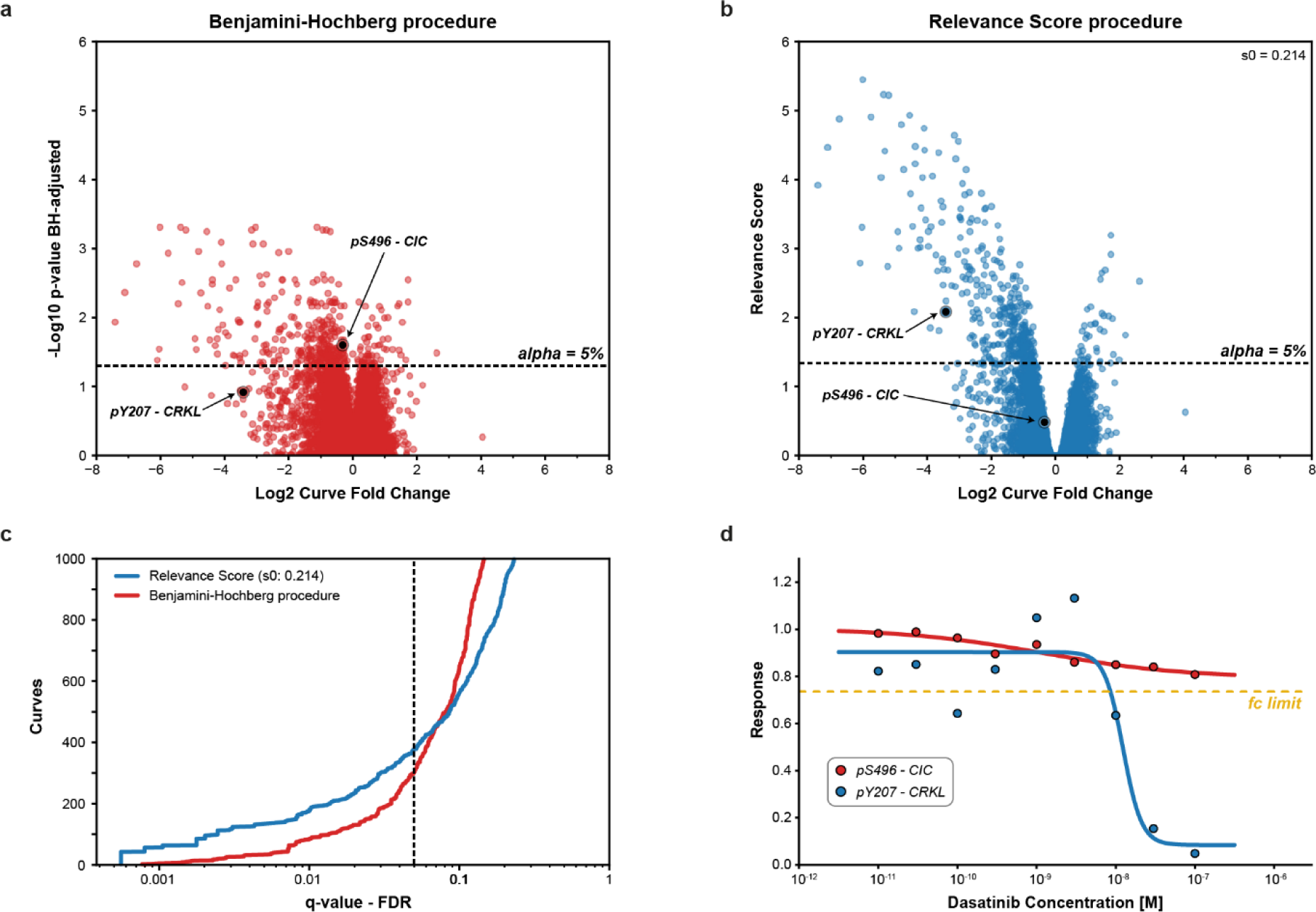
Relevance asymptotes show better performance than multiple testing correction. DecryptM data (dose-response phosphorylation data) of replicates of Dasatinib-treated K562 cells was processed by CurveCurator. In this biological experiment, only a few phosphorylation sites (p-sites; low 100s) were expected to change, and the majority of p-sites (>10,000) was expected to be unresponsive to the treatment. This scenario requires very high statistical sensitivity. Two phosphorylation sites are highlighted to illustrate the difference between multiple testing corrections and the relevance score approach. pY207 (phospho-tyrosine modification at position 207) on CRKL is a known substrate of the tumor-driving fusion kinase BCR-ABL and can be regarded as a true responsive site of Dasatinib treatment. pS496 (phospho-serine modification at position 496) on CIC neither has a functional annotation nor an established kinase-substrate relationship. As BCR-ABL is a phospho-tyrosine kinase, pS496 cannot be a direct substrate of BCR-ABL. **a)** P-values were multiple testing corrected using the Benjamini-Hochberg (BH) procedure, which only uses the p-value dimension for FDR correction. Here, the CIC pS-site but not the CRKL pY-site would be called significantly regulated. **b)** P-Values were transformed into the relevance scores using an alpha value of 5% and a log_2_ fold change limit of 0.45, resulting in an s_0_ factor of 0.214. Now, the CRKL pY-site but not the CIC pS-site is correctly identified as regulated. **c)** Comparing BH q-values with relevance score q-values indicates that the latter can identify more significant dose-response curves in the lower FDR range. At high FDRs, BH finds more regulated curves compared to the relevance score because the fold-change limit prevents those biologically less relevant but statistically significant curves from being called regulated. **d)** Experimentally determined dose-response curves of the selected examples pY207 on CRKL (red) and pS496 on CIC (blue). The log_2_ fold change limit of 0.45 is indicated by the yellow line. It is evident that pY207 on CRKL is drug-regulated and pS496 o

